# Tax induces the recruitment of NF-kB to unintegrated HIV-1 DNA to rescue viral gene expression and replication

**DOI:** 10.1101/2021.02.16.431555

**Authors:** Ishak D. Irwan, Bryan R. Cullen

## Abstract

We have previously reported that the normally essential step of integration of the HIV-1 proviral DNA intermediate into the host cell genome becomes dispensable in T cells that express the Human T cell leukemia virus 1 (HTLV-1) Tax protein. The rescue of integrase (IN) deficient HIV-1 replication by Tax results from the strong activation of transcription from the long terminal repeat (LTR) promoter on episomal HIV-1 DNA, an effect that is closely correlated with the recruitment of activating epigenetic marks, such as H3Ac, and depletion of repressive epigenetic marks, such as H3K9me3, from chromatinized unintegrated proviruses. In addition, activation of transcription from unintegrated HIV-1 DNA coincides with the recruitment of NF-kB to the two NF-kB binding sites found in the HIV-1 LTR enhancer. Here we report that the recruitment of NF-kB to unintegrated viral DNA precedes, and is a prerequisite for, Tax-induced changes in epigenetic marks, so that an IN-HIV-1 mutant lacking both LTR NF-kB sites is entirely non-responsive to Tax and fails to undergo the epigenetic changes listed above. We also report that heterologous promoters introduced into IN-HIV-1-based vectors are transcriptionally active even in the absence of Tax. Finally, we failed to reproduce a recent report arguing that heterologous promoters introduced into IN-vectors based on HIV-1 are more active if the HIV-1 promoter and enhancer, located in the LTR U3 region, are deleted, in a so-called self inactivating or SIN lentivector design.

**Importance:** Integrase-deficient expression vectors based on HIV-1 are becoming increasingly popular as tools for gene therapy *in vivo* due to their inability to cause insertional mutagenesis. However, many IN-lentiviral vectors are able to achieve only low levels of gene expression and methods to increase this low level have not been extensively explored. Here we analyze how the HTLV-1 Tax protein is able to rescue the replication of IN-HIV-1 in T cells and describe IN-lentiviral vectors that are able to express a heterologous gene effectively.

## Introduction

Integration of the linear DNA proviral intermediate into host cell DNA by the viral integrase (IN) enzyme is not only a key step in the life cycle of all retroviral species examined so far but also essential for effective retroviral gene expression (1,2). As a result, the HIV-1 IN protein has emerged as a major target for antiviral drugs, such as raltegravir and dolutegravir, that function as IN strand transfer inhibitors and thereby effectively prevent integration (3–5). Importantly, these drugs not only block HIV-1 replication and gene expression in dividing cells, such as T cells, but also in non-dividing cells, such as macrophages, thereby demonstrating that loss of unintegrated proviral DNA during cell division, though this clearly does occur, is at least not the only inhibitory process that proviral integration has evolved to avoid (6–9). In fact, it is now clear that the inability of unintegrated proviral DNA to support effective retroviral gene expression and replication is due to the epigenetic silencing of episomal retroviral DNA (10–14). This involves the rapid recruitment of repressive epigenetic marks, such as H3K9me3, to chromatinized retroviral episomes shortly after nuclear entry and also correlates with the inability of unintegrated retroviral DNA to recruit the activating epigenetic marks, such as H3Ac, that are abundant on actively transcribed, integrated proviruses.

While proviral activation is therefore normally an essential step in the HIV-1 life cycle, we recently demonstrated that HIV-1 IN function is dispensable in T cells that express an inducible form of the Human T cell leukemia virus 1 (HTLV-1) Tax protein (13,15). Specifically, induction of Tax expression in the CD4+ T cell line CEM-SS led to a high level of transcription from the HIV-1 long terminal repeat (LTR) promoter present on unintegrated viral DNA that was sufficient to support the robust replication and spread of HIV-1 in culture. The Tax-mediated activation of the HIV-1 LTR coincided with not only the recruitment of activating epigenetic marks, such as H3Ac, to unintegrated HIV-1 DNA at levels that actually exceeded the level of H3Ac detected on integrated proviruses but also prevented the acquisition of repressive epigenetic marks, such as H3K9me3.

Because integration of retroviral proviruses into the host cell genome has the potential to cause insertional mutagenesis, which could potentially lead to oncogenic transformation (16,17), there has been considerable interest in the question of whether one could design lentiviral vectors based on HIV-1 or other lentiviruses that remain able to express significant levels of a heterologous gene of interest in the absence of IN function (18,19). Initially, several groups reported that non-integrating vectors based on HIV-1, feline immunodeficiency virus (FIV) or equine infectious anemia virus (EIAV) produced negligible levels of an encoded indicator gene, >100-fold lower that seen with the matched IN+ control vector (20–23). However, more recently reasonable levels of gene expression by IN-vectors have been reported by several groups, though no clear understanding of why some designs appear more effective than others has been presented (24–27). While it has been reported that deletion of almost the entire U3 region in the 3’ LTR, to generate a so-called self-inactivating or SIN lentivector, increased expression of the encoded green fluorescent protein (GFP) by ~3 fold, this level still remained ~7 fold lower than seen with the matched IN+ vector (28).

In this manuscript, we have sought to further define why an HIV-1 mutant lacking a functional IN protein is replication defective in normal T cells but able to replicate in the presence of the HTLV-1 Tax protein. We now report that this effect is due to the activation of NF-kB by Tax, resulting in the effective recruitment of the NF-kB subunits Rel A and Rel B to the two NF-kB binding sites located in the HIV-1 LTR U3 region on unintegrated proviral DNA. This in turn modifies the chromatin marks recruited to these retroviral episomes, leading to a highly significant increase in the activating H3Ac epigenetic mark, and a marked reduction in the repressive H3K9me3 epigenetic mark. Finally, we report the efficient expression of an indicator gene product from several HIV-1-based IN-vectors even in the absence of Tax and find that, in our hands, the SIN vector design is not more active.

## Results

We have previously reported that expression of the HTLV-1 Tax protein in human T cells, in particular in the CD4+ human T cell lines C8166 and CEM-SS, rescues gene expression from unintegrated DNA episomes derived from an IN-HIV-1 mutant and allows their robust replication despite their lack of integrase function (13). We further reported that this effect correlated with both the recruitment of the NF-kB subunits Rel A and Rel B to the two NF-kB sites located in the HIV-1 LTR enhancer and with the addition of the activating epigenetic marks H3Ac and H3K4me3, and the loss of the repressive epigenetic mark H3K9me3, from chromatin bound to unintegrated HIV-1 episomes. However, our previous work only looked at the epigenetic status of unintegrated viral DNA late after infection and therefore we could not determine whether the observed rescue of HIV-1 gene expression was due to the observed epigenetic changes leading to the effective recruitment of NF-kB to unintegrated HIV-1 DNA, or *vice versa*. We reasoned that it might be possible to resolve this question by addressing the temporal order underlying the recruitment of the NF-kB subunits, relative to the addition of epigenetic marks, to unintegrated HIV-1 DNA. For this purpose, we infected Tax-inducible CEM-SS cells with IN+ or IN-forms of the previously described replication competent indicator virus NL-NLuc (29) in the presence or absence of doxycycline (Dox) and then harvested the infected cells at 16, 19, 21 or 24 hours post-infection (hpi). We then performed chromatin immunoprecipitation linked to qPCR (ChIP-qPCR) to quantify the level of Rel A, Rel B, H3Ac and H3K9me3 present on proviral DNA. As shown in Fig. 1A and 1B, analysis of the recruitment of Rel A and Rel B to the LTR on unintegrated HIV-1 DNA revealed a highly significant (p<0.0001) boost in the recruitment of both proteins to the HIV-1 LTR in Tax expressing CEM-SS cells, relative to uninduced CEM-SS cells, as early as 16 hpi, the earliest time point examined, and this effect was even clearer at subsequent times. In contrast, as shown in Figs. 1C and 1D, we did not detect any increase in the recruitment of H3Ac to HIV-1 DNA-bound chromatin, or loss of H3K9me3 marks from viral chromatin, at 16 hpi though this effect did become significant (p<0.001) at 19 hpi and was even more pronounced at later times, as also previously reported.

**Figure 1:**
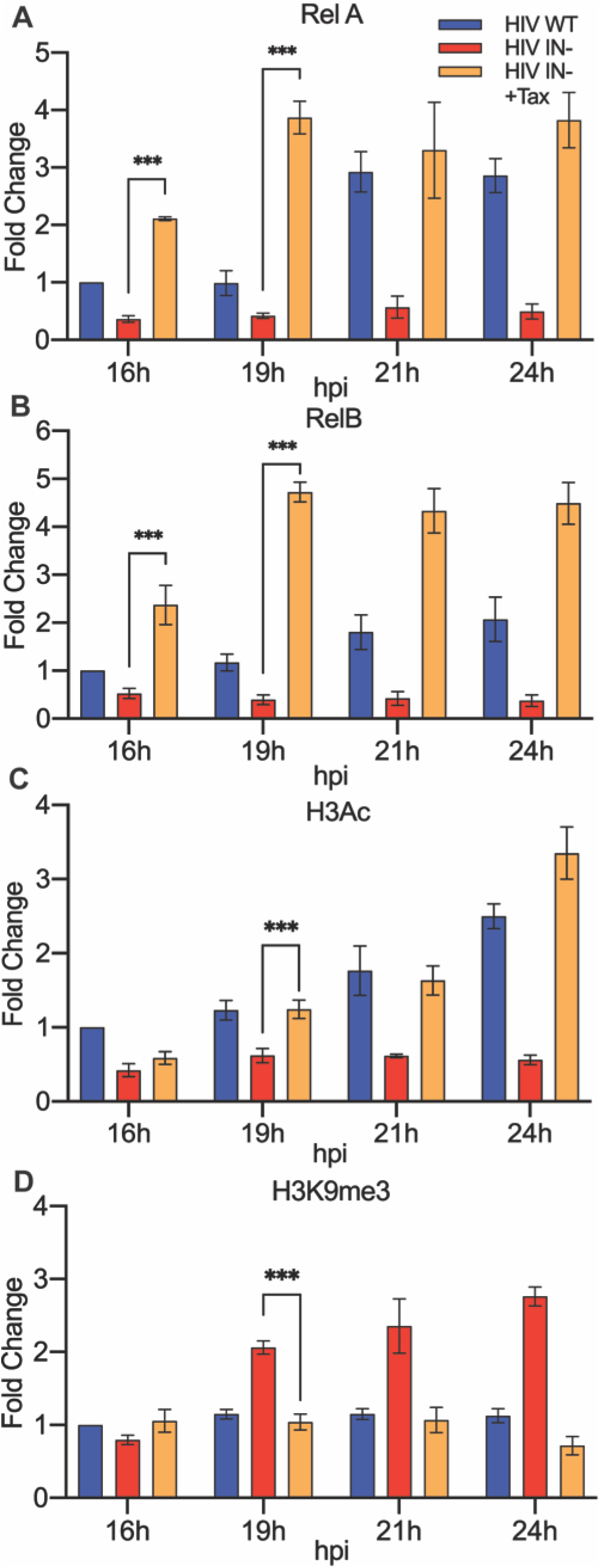
NF-kB subunits RelA and RelB are recruited to the HIV-1 LTR prior to activating or repressive epigenetic histone modifications. ChIP-qPCR detection of (A) RelA/p65, (B) Rel B, (C) acetylated histone H3 (H3Ac), or (D) tri-methylated histone 3 lysine 9 (H3K9me3) loaded on to the HIV-1 LTR at the indicated hours post-infection (hpi) in CEM-SS cells expressing or not expressing the HTLV-1 Tax protein. Ct values were corrected for total H3 detected on the GAPDH gene and normalized to the WT HIV-1 infection at 16 hpi, which was set to 1. n=3, ± SD. ***p<0.001, calculated by 2-Way ANOVA with Tukey’s multiple comparison test.

The data presented in Fig. 1 reveal that Tax-induced Rel A and Rel B are recruited to unintegrated HIV-1 proviruses before any Tax-induced changes in either activating (H3Ac) and repressive (H3K9me3) chromatin marks can be detected. However, these data do not address whether the Tax-induced activation and recruitment of NF-kB to viral DNA is necessary for the rescue of IN-HIV-1 replication. To address this question, we generated derivatives of the previously described IN-form of the indicator virus NL-NLuc (13) in which either one LTR NF-kB site (1Δ) or both NF-kB sites (2Δ) were mutationally inactivated. We then asked whether loss of NF-kB sites affected the rescue of this IN-virus by Tax. In this experiment, we infected CEM-SS cells bearing the Tet-inducible *tax* gene with IN+ NL-NLuc, with IN-NL-NLuc, or with the new 1Δ or 2Δ NF-kB mutants of IN-NL-NLuc. We then measured the level of viable cells in each culture over time (Fig 2A), the level of expression of the virally encoded NLuc indicator (Fig. 2B), the level of viral DNA (Fig. 2C) and the level of total viral RNA (Fig. 2D) at up to 7 days post-infection (dpi). As shown in Fig. 2A, and as previously reported, we observed that infection of CEM-SS cells with the IN-NL-NLuc derivative bearing an intact LTR did not reduce cellular viability, relative to the uninfected culture, in the absence of but induced the death of almost the entire CEM-SS culture by seven days post-infection (dpi) if Tax expression was induced by Dox addition. Interestingly, while the 1Δ mutant of the IN-NL-NLuc virus also induced a strong cytopathic effect, the 2Δ mutant phenocopied the lack of cell killing seen with the parental NL-NLuc virus in the absence of Tax (Fig. 2A). The lack of cytopathic effect seen with the 2Δ mutant of NL-NLuc correlated with an almost complete lack of viral gene expression, as measured by NLuc activity (Fig. 2B). In contrast, the 1Δ mutant, which retains a single NF-kB site, gave rise to a lower level of NLuc activity than the parental IN-NL-NLuc virus, which retains both NF-kB sites, that was nevertheless readily detectable (Fig. 2B). Similarly, the 2Δ mutant failed to generate a spreading infection in Tax-expressing CEM-SS cells, as measured by our inability to measure any increase in either viral DNA or viral RNA expression over time (Figs. 2C and D). The 1Δ mutant was again intermediate in phenotype between the 2Δ mutant and the parental NL-NLuc vector in that it remained able to spread through the Tax-expressing CEM-SS culture, albeit more slowly than NL-NLuc, as revealed by increasing levels of both HIV-1 DNA and HIV-1 RNA expression (Figs. 2C and D).

**Figure 2:**
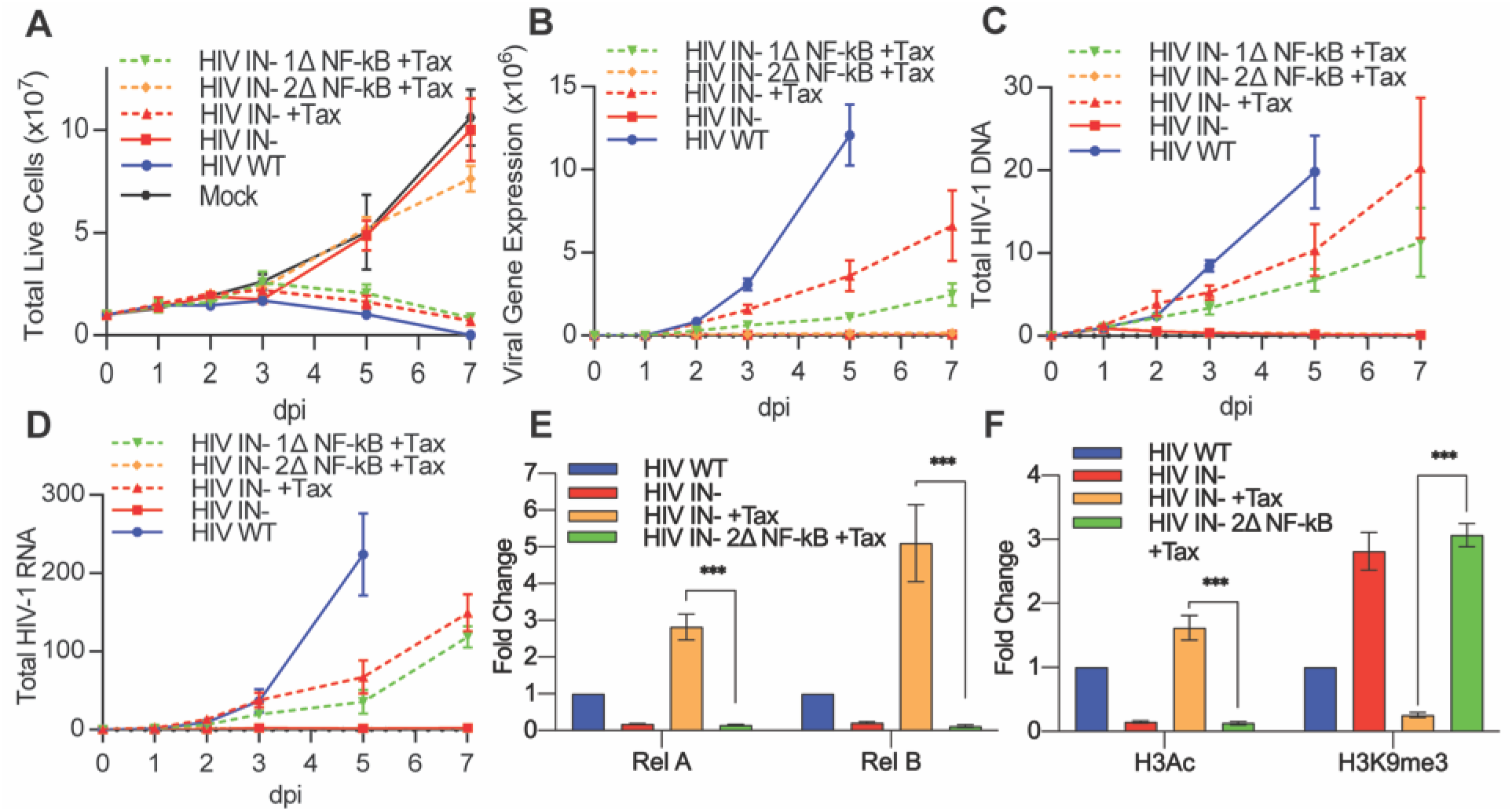
At least one HIV-1 LTR NF-kB binding site is required for the rescue of IN-HIV-1 replication by Tax. Clonal CEM-SS T cell lines with Tet-inducible Tax were infected with WT or IN-NL-NLuc with either neither, one, or both of the NF-kB sites mutated. (A) Live cell counts (B) Virally encoded NLuc expression (C) total HIV-1 DNA measured by qPCR or (D) HIV-1 RNA expression measured by qRT-PCR were quantified at 1, 2, 3, 5 and 7 days post-infection (dpi) post-infection. The WT culture was devoid of live cells at 7 dpi. DNA and RNA levels were normalized to WT HIV-1 infected cells in the absence of Tax at 1 dpi, which was set to 1. The epigenetic state of the viral LTR promoter was characterized by quantifying (E) the bound NF-kB sub-units Rel A and Rel B and (F) the activating H3Ac and repressive H3K9me3 histone modification at 2 dpi using ChIP-qPCR. n=3, ±SD. ***p<0.001, calculated by 2-Way ANOVA with Tukey’s multiple comparison test.

The data presented in Figs. 2A-D demonstrate that the loss of both NF-kB sites present in the HIV-1 LTR prevents the rescue of viral gene expression by Tax seen when both or even only one LTR NF-kB site is retained in the LTR of an IN-HIV-1 mutant, a result that is consistent with the known ability of Tax to activate NF-kB in expressing cells (15). These data could suggest that recruitment of NF-kB is required for the subsequent epigenetic changes, that is increased H3Ac marks and reduced H3K9me3 marks, seen on unintegrated HIV-1 proviruses after rescue by Tax induction (Figs. 1C and D). Alternatively, it could be argued that these epigenetic changes are still induced by Tax but, in the absence of NF-kB recruitment, are insufficient to rescue transcription of unintegrated HIV-1 DNA. To resolve this question, we performed CHiP-qPCR on IN-NL-NLuc either bearing a WT LTR promoter or the 2Δ LTR lacking both NF-kB sites. As expected, in the absence of Tax, we observed almost no detectable recruitment of either the Rel A or Rel B NF-kB subunit to the intact viral LTR on unintegrated viral DNA (Fig. 2E) and similarly, in the absence of Tax, the chromatin added to unintegrated HIV-1 episomes was modified by addition of high levels of the repressive H3K9me3 mark while the activating H3Ac mark was essentially absent (Fig. 2F). In contrast, in the presence of Tax, both Rel A and Rel B were effectively recruited to the intact LTR promoter present on unintegrated HIV-1 DNA (Fig. 2E) and we now saw efficient recruitment of the activating H3Ac mark, and very little addition of the repressive H3K9me3 mark (Fig. 2F), as expected. Importantly, however, when the 2Δ mutant of IN-NL-NLuc was examined in the presence of Tax it phenocopied the pattern observed for IN-NL-NLuc bearing the WT LTR in the absence of Tax. That is, while the 2Δ indicator virus mutant failed to recruit either the Rel A or Rel B NF-kB subunit, as expected (Fig. 2E), it was heavily modified by addition of the repressive H3K9me3 mark to chromatinized viral DNA while remaining devoid of the activating H3Ac marks seen on the unintegrated WT LTR in the presence of Tax (Fig. 2F). Thus, these data argue that recruitment of the NF-kB subunits Rel A and/or Rel B to the viral LTR present on unintegrated HIV-1 DNA is a prerequisite for the subsequent recruitment of the epigenetic marks that correlate with active transcription of a DNA template.

While there is considerable evidence indicating that integration of the retroviral proviral intermediate into the host cell genome is an essential step in the retroviral life cycle (1,2), and demonstrating that this is largely due to the efficient epigenetic silencing of unintegrated retroviral DNA (10–14), there are also a number of reports documenting the effective expression of cDNAs encoding a gene of interest from some, but not all, integrase deficient lentiviral vectors bearing inserted heterologous enhancer/promoter combinations (24–27). To our knowledge, the issue of why some, but not all, lentiviral vectors are able to escape epigenetic silencing has not been resolved, though it has been suggested that deletion of the retroviral promoter/enhancer located in the LTR U3 region, in so-called self-inactivating or SIN lentiviral vectors, can promote the function of inserted heterologous promoters (28).

To address this question, we generated a set of vectors from the WT and IN-NL-NLuc vectors in which we first deleted almost the entire U3 region, from −18 to −418, in the 3’ LTR, while leaving *cis*-acting sequences required for reverse transcription and integration intact, to generate SIN vectors (30). We then inserted either the LTR enhancer/promoter from Rous Sarcoma Virus (RSV) (31) or the highly active cytomegalovirus immediate early enhancer/promoter (CMV) (32), 5’ to the NLuc indicator gene (Fig. 3A). These vectors were then packaged and used to infect Tax-inducible CEM-SS cells in the presence and absence of Dox. At 48 hpi, the cells were lysed and the level of virus-encoded NLuc determined. As may be seen in Fig. 3B, the level of NLuc expression induced upon infection with lentiviral vectors containing internal heterologous promoters was comparable to that seen with the parental NL-NLuc vector and, unlike NL-NLuc, was only modestly inhibited, by <3-fold, when the integrase gene was mutationally inactivated. While expression of Tax did not affect the level of NLuc expression from either the IN+ or IN-forms of NL-SIN-RSV-NLuc, Tax did exert a significant positive effect on expression from the CMV promoter, which contains NF-kB binding sites (34), in both the IN+ and IN-version of NL-SIN-CMV-NLuc (Fig. 3B).

**Figure 3:**
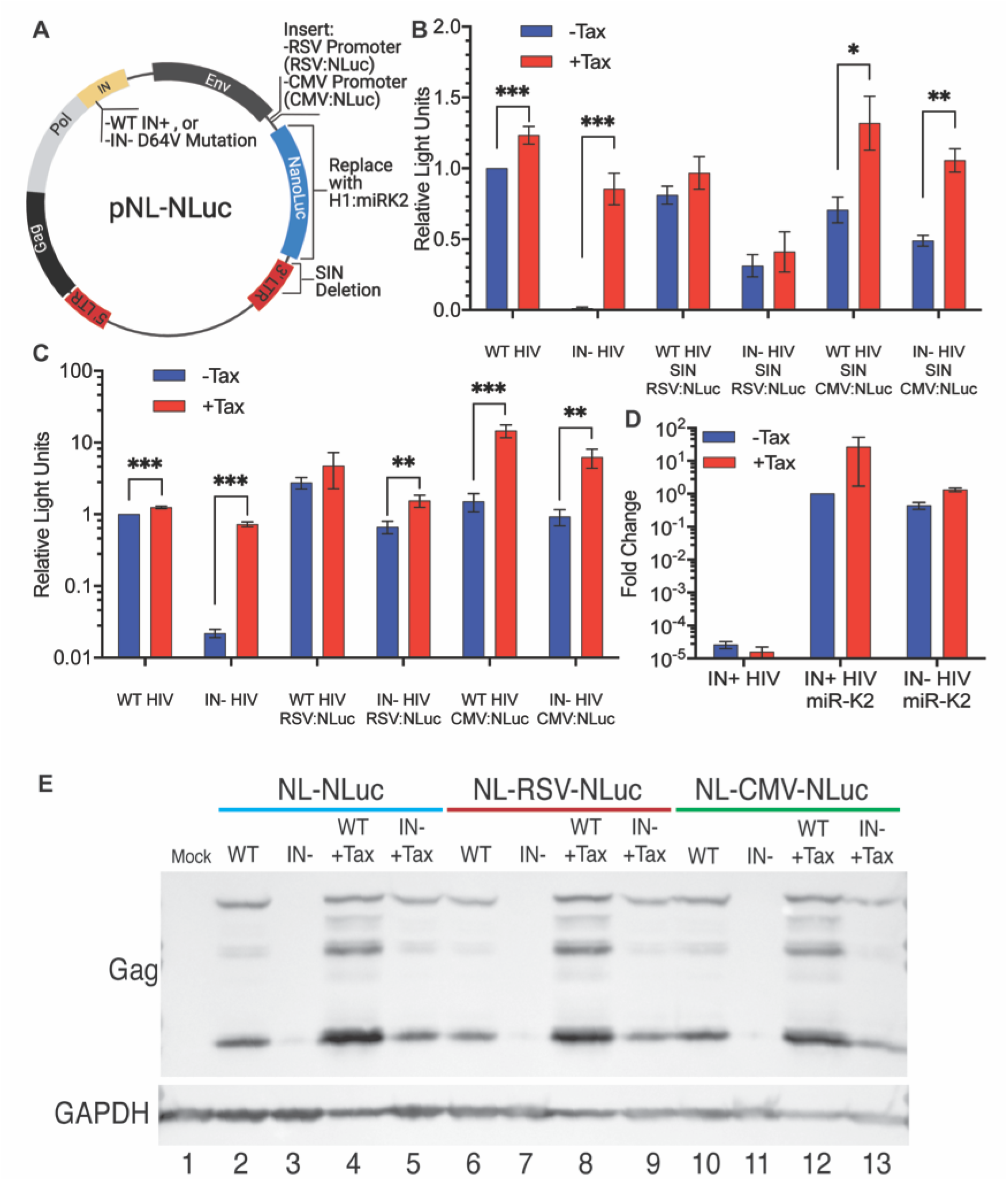
Unlike the HIV-1 LTR, heterologous internal Pol II and Pol III promoters are not silenced on unintegrated HIV-1 proviruses. (A) Schematic of pNL-NLuc and its derivatives. (B) CEM-SS cells *±* Tax were infected with IN+ or IN-HIV-1 variants expressing NLuc from the viral LTR, or with self-inactivating (SIN) derivatives lacking the LTR U3 region from −18 to −418, including the HIV-1 promoter and enhancer. Instead these viruses express NLuc from an internal RSV LTR promoter (that has no NF-kB binding sites), or an internal CMV promoter (that has NF-kB binding sites). Nevirapine was added at 16 hpi to prevent virus spread, and cells were harvested for NLuc assays at 48 hpi. NLuc activity was normalized to WT NL-NLuc in the absence of Tax, which was set to 1.(C) As in panel B but using a non-SIN vector design. (D) CEM-SS *±* Tax cells were infected with WT or IN-virus expressing KSHV miR-K2 from the Pol III-dependent H1 promoter. Nevirapine was added to the cells at 16 hpi, and the cells harvested for quantification of miR-K2 by qRT-PCR at 48 hpi. Endogenous U6 RNA served as an internal control and data were normalized to WT minus Tax, which was set to 1. n=3, *±* SD. (E) HIV-1 Gag expression levels in CEM-SS cells +/-Tax infected with the indicated WT or IN-viruses. Endogenous GAPDH was used as a loading control. *p<0.05, **p<0.01, *** p<0.001, calculated using an unpaired Holm-Sidak multiple comparison t-test.

As noted above, the SIN design has been reported to enhance the level of gene expression from heterologous promoters inserted into IN-retroviral vectors (28), possibly due to the loss of the potential for transcriptional interference inherent in this design (33). To test whether this is indeed the case, we generated a set of four IN+ or IN-vectors, bearing either the RSV or CMV enhancer/promoter driving NLuc expression, that differed from the vectors analyzed in Fig. 3B only in that the HIV-1 LTR was now left intact. As shown in Fig 3C, these vectors in fact expressed up to 10-fold higher levels of NLuc that the similar SIN vectors. This higher level of NLuc expression was slightly more subject to inhibition upon loss of IN function than seen with the SIN design and was also more responsive to the HTLV-1 Tax protein. This latter effect was again particularly prominent with vectors bearing the CMV enhancer/promoter which, unlike the RSV LTR, contains NF-kB binding sites (34). Nevertheless, overall it was clear that the tested lentiviral vectors, bearing internal heterologous promoters, were all transcriptionally active regardless of IN activity and that the SIN design did not increase expression from unintegrated lentiviral vector DNA, and in fact seemed to modestly inhibit NLuc expression.

Finally, while the data presented in Fig. 3A and B reveal that a heterologous Pol II-dependent promoter inserted into a lentiviral vector lacking integrase function is still essentially fully active, they do not address whether other types of promoters, for example promoters that utilize Pol III, would also be functional on a non-integrated proviral DNA template. This could be important if, for example, one wished to express a microRNA (miRNA) from a non-integrating retroviral vector. To tes**t** this, we therefore inserted a previously described expression cassette containing the H1 Pol III promoter linked to an RNA hairpin encoding the miR-K2 miRNA expressed by Kaposi’s Sarcoma Herpesvirus (KSHV) (35). As may be seen in Fig. 3C, this vector expressed high levels of mature miR-K2, as measured by qRT-PCR, regardless of whether an IN+ or IN-vector design was used. Unexpectedly, the expression of miR-K2 appeared to be enhanced, especially in the IN+ vector, by induction of Tax expression.

The CMV immediate early promoter contains an exceptionally active enhancer (32), and, given that both the CMV and RSV promoters were active in the context of the NL-NLuc vectors bearing intact LTRs, we wondered if the CMV or RSV promoter/enhancer might be able to rescue expression from the 5’ HIV-1 LTR in *cis* when integrase was mutationally inactivated. If so, this should result in the expression of the HIV-1 Gag protein. To test this, we infected CEM-SS cells with NL-NLuc, with NL-RSV-NLuc and with NL-CMV-NLuc in the presence and absence of an intact integrase gene and in the presence or absence of Dox, to induce Tax expression, As expected, and as previously reported (13), the IN-form of NL-NLuc failed to express Gag unless Tax expression had been induced (Fig. 3E, compare lanes 3 and 5). Similarly, for both NL-RSV-NLuc and NL-CMV-NLuc, we failed to detect Gag expression from the IN-form of this vector unless Tax expression was induced (Fig. 3D, lanes 7, 9, 11 and 13). We therefore conclude that, even though both the RSV LTR and CMV promoter/enhancer remain essentially fully active when inserted into a lentiviral vector lacking integrase function, they are nevertheless both unable to prevent the epigenetic silencing of the immediately adjacent HIV-1 LTR promoter.

## Discussion

Previously, we reported that expression of the HTLV-1 Tax protein in human CD4+ T cells is able to rescue the transcription and robust replication of an integrase-deficient HIV-1 mutant, bearing the D64V IN mutation, in the absence of any detectable proviral DNA integration (13). This rescue correlated with both the recruitment of activated NF-kB to the HIV-1 LTR on unintegrated DNA and with the acquisition of high levels of the activating epigenetic mark H3Ac, and loss of the repressive mark H3K9me3. While our previous research did not address whether the rescue of HIV-1 gene expression by Tax was primarily due to NF-kB activation and recruitment or, instead, due to changes in the epigenetic status of the HIV-1 LTR promoter, we did note that a Tax mutant that had selectively lost the ability to activate NF-kB also failed to rescue IN-HIV-1 replication (13).

Here, we extend this earlier work by demonstrating that recruitment of the NF-kB subunits Rel A and Rel B to the HIV-1 LTR on unintegrated viral DNA precedes the acquisition of activating H3Ac marks, and loss of repressive H3K9me3 marks, by the same viral DNA. Moreover, we observed that mutational inactivation of both NF-kB sites on the HIV-1 LTR not only blocks the recruitment of NF-kB subunits to unintegrated viral DNA but also entirely blocks the epigenetic changes, noted above, that mark DNA as being transcriptionally active. We have therefore demonstrated that activation of NF-kB by Tax (36), leading to its recruitment to unintegrated HIV-1 DNA, is the key step that then results in the epigenetic changes that correlate with the active transcription and rescue of IN-HIV-1 mutants.

In addition to addressing the effect of Tax on IN-full length HIV-1, we also looked at how loss of IN function, and activation of Tax expression, affects gene expression from a heterologous reporter cassette inserted into a SIN configuration of the HIV-1 genome. As shown in Fig. 3B, we observed a high level of expression from inserted RSV- and CMV-derived enhancer/promoter combinations driving expression of the NLuc indicator gene and this high level of expression was, unexpectedly, only minimally impacted by loss of IN function. This high level of NLuc expression, in both the presence and absence of IN function, was also observed when an HIV-1 based vector retaining a fully intact LTR promoter was analyzed, thus arguing that, at least in this vector design, the HIV-1 LTR does not exert any inhibitory effect in *cis*. To confirm that the intact HIV-1 LTR present in these vectors was indeed active, we were able to demonstrate high levels of HIV-1 Gag protein expression from the IN+ version of the NL-RSV-NLuc and NL-CMV-NLuc vectors as well as from the IN-version, but only in the presence of Tax in the latter case. Therefore, while both the RSV LTR and CMV immediate early promoter-enhancer are highly active in the context of unintegrated HIV-1 DNA, they are both unable to activate the HIV-1 LTR promoter in *cis*.

## Materials and Methods

### Cells and Cell Culture Conditions

The cell lines used in this study are HEK293T (293T), a human kidney epithelial cell line of female origin, and CEM-SS, a human CD4+ T cell line of female origin. CEM-SS cells transduced with a lentiviral vector bearing a Tet-inducible promoter that controls the expression of the HTLV-1 Tax protein, have been previously described (13). All cells were cultured at 37C with 5% CO2. 293T cells were cultured in Dulbecco’s Modified Eagle Medium (DMEM) supplemented with 10% fetal bovine serum (FBS) and Antibiotic-Antimycotic. CEM-SS cells were cultured in Roswell Park Memorial Institute (RPMI) medium supplemented with 10% FBS and Antibiotic-Antimycotic.

### HIV-1 Production

In experiments that measure the level of viral NLuc expression in cells infected with replication competent HIV-1, we used a previously described replication competent NLuc reporter virus (pNL-NLuc) where the viral *nef* gene in NL4-3 was replaced with the *NLuc* gene (29). The 1Δ- and 2Δ NF-κB mutant viruses were created by mutating either one or both of the NF-κB binding sites in the HIV-1 LTR U3 region, respectively, by using overlap-extension PCR to change the NF-κB binding site from the functional 5’-GGGACTTTCC-3’ to a non-functional *5’-GGGACTGGAT-3’* sequence.

HIV-1 expressing miR-K2 (pNL-miR-K2) was created by PCR amplification of an H1 promoter-driven miR-K2 expression cassette from pSUPER-miR-K2 (35), followed by cloning into pNL4-3 using Not I and Xho I. HIV-1 SIN viruses expressing NLuc from an RSV or CMV-derived promoter were created by first cloning the promoters into the Not I site 5’ of the *NLuc* gene in pNL-NLuc (Fig. 3A), then replacing the 3’LTR with the corresponding SIN LTR from the pNL-SIN-CMV-RLuc plasmid described previously (37), which bears a deletion extending from −18 to −418 in U3.

All viruses express either WT integrase (IN), or IN bearing a point mutation, D64V, that blocks IN function (38). Plasmids expressing the NL-NLuc provirus, or the relevant mutant constructs, were transfected into 293T cells using polyethylenimine (PEI). At 24h post-transfection, the spent media were replaced with fresh media. At 72 h post-transfection, supernatant media were harvested and passed through a 0.45um filter to remove cell debris. HIV-1-containing supernatants were normalized using a p24 ELISA before being used to infect target cells.

### ChIP-qPCR

Tet-inducible HTLV-1 Tax CEM-SS cells were cultured in 10ml of RPMI media (at 10^6^ cells/ml) and infected with WT or IN-HIV-1 virus in the presence or absence of 0.5ug/ml Dox to induce Tax production. Prior to infection, viral supernatants were incubated with 5U/ml of DNAse I at 37°C for 1 hour to remove residual plasmid contamination.

Cells were harvested at the indicated times post-infection, rinsed twice with 1x PBS and cross-linked with 1% formaldehyde for 20 min at 25 °C, before being quenched in 0.125 M glycine for 5 min. The rinsed cells were then lysed and subjected to chromatin immunoprecipitation (ChIP) as previously described (13) to pull down the DNA associated with the indicated NF-κB subunits or modified histones. DNA was then purified using DNA cleanup columns (Zymo Research), digested with DpnI (NEB) to remove any residual plasmid contamination, then used for qPCR analysis using primers that amplify U5-R on HIV-1 in a SYBR green master mix (ThermoFisher). ΔΔCt was calculated relative to total histone H3 levels and expressed as a fold change relative to CEM-SS cells not expressing Tax and infected with WT HIV-1.

### Luciferase Assays

Cells were washed three times in PBS, lysed in Passive Lysis Buffer (Promega) and assayed for NLuc activity using the Nano-Glo Luciferase Assay on a Lumat LB9507 luminometer (Bertold Technologies).

### Quantifying Changes in HIV-1 Replication and Viral Gene Expression

10^7^ Tet-inducible Tax CEM-SS cells were suspended in 10ml of RPMI medium and infected with NLuc reporter viruses in the presence or absence of 0.5ug/ml Dox to induce Tax expression. Viral supernatants were normalized by p24 and pretreated with 5 U/mL DNase I for 1 h at 37 °C to remove residual plasmid DNA. All IN-HIV-1 infections were supplemented with raltegravir (20uM) to prevent any revertant mutations. Live cells were quantified and 10^6^ live cells harvested on days 1, 2, 3, 5, and 7 and equally split into 3 aliquots to assay NLuc or to extract and analyze viral DNA and RNA.

For DNA analysis, cells were spun down and washed three times in ice cold PBS and the cells incubated with DpnI (NEB) to remove any residual plasmid contamination. DNA was then extracted using Quick-DNA Miniprep Plus columns (Zymo Research).

For RNA analysis, cells were lysed in TRIzol (ThermoFisher) and RNA harvested according to the manufacturer’s instructions, and DNaseI treated for 2h. The DNaseI was then heat inactivated, and the RNA converted to cDNA using a High Capacity cDNA Reverse Transcription kit (Applied Biosystems).

All qPCR assays were performed in triplicate in a QuantStudio 6 Pro real-time PCR system according to the manufacturer’s instructions. HIV-1 DNA/cDNA was amplified with a custom total HIV-1 TaqMan probe/primer set that amplifies the U5-gag region of HIV-1(39). β-Actin DNA was PCR amplified using a premade TaqMan probe/primer set (ThermoFisher, Cat #431182), while β-Actin cDNA in RNA analyses was quantified using a separate Taqman probe/primer set that amplifies across a splice junction (FP: 5’-CAATGAAGATCAAG ATCATTGC-3’, RP: 5’-AAGCATTTGCGGTGGAC-3’, probe: 5’-FAM-CCACCTTCCAGCAGATGTGGATCAGCAAG-TAMRA-3’). Relative quantification of HIV-1 DNA/RNA levels was performed using the ΔΔCT method by normalizing total HIV-1 DNA/RNA to the β-Actin internal control (40).

### Quantification of miR-K2

Tet-inducible HTLV-1 Tax CEM-SS cells were infected with IN+ or IN-pNL-miR-K2 in the presence or absence of 0.5ug/ml Dox. Nevirapine (Sigma) was added to the cells at 16hpi, and the cells harvested for RNA analysis using Trizol (ThermoFisher). RNA was diluted to 2ng/ul then reverse transcribed with either a specific miR-K2 primer or a U6 RNA internal control primer (Applied Biosystems) using a MicroRNA Reverse Transcription kit (Applied Biosystems). The converted cDNA was then quantified by Taqman qPCR small RNA assays that target either miR-K2 or U6. All qPCR assays were performed in a QuantStudio 6 Pro real-time PCR machine. Quantification was performed by correcting the observed miR-K2 level to the U6 internal control. Corrected miR-K2 expression levels were then normalized to the level of miR-K2 detected in cells infected with IN+ pNL-miR-K2 in the absence of Tax, which was set to 1.

### Western blot for HIV-1 Gag

Cells were lysed in Laemmli buffer, sonicated and denatured at 95°C for 10 min. Lysates were run on a 4–20% SDS-polyacrylamide gel (Bio-Rad), transferred to a nitrocellulose membrane and then blocked in 5% milk in PBS + 0.1% Tween (PBS-T). Membranes were incubated with a p24-specific monoclonal antibody (#24-3) (AIDS Reagent, ARP: 6458), or a glyceraldehyde 3-phosphate dehydrogenase (GAPDH) antibody (Proteintech, 60004-1), followed by a secondary anti-mouse HRP antibody (Sigma, A9044) in 5% milk in PBS-T for 1 h each and then washed in PBS-T. The membranes were incubated with a luminol-based enhanced chemiluminescent (ECL) substrate (Advansta) and signals visualized using GeneSnap (Syngene).

## Acknowledgements

This research was supported in part by NIH grant R21-AI157616 to B.R.C., who is also supported through the Center for HIV RNA Studies (CRNA, U54-AI150470). This research received infrastructure support from the Duke University CFAR (P30-AI064518). The following reagent was obtained through the NIH AIDS Reagent Program, Division of AIDS, NIAID, NIH: p24 Gag Monoclonal (#24-3) from Michael Malim.

